# Vocal Tract Disparity and Potential Implications for Speaker Recognition

**DOI:** 10.64898/2026.08.24.746615

**Authors:** Tiena Danner, Valeriia Vyshnevetska, Daniel Friedrichs, Steven Moran

**Affiliations:** Institute of Biology, University of Neuchâtel, 2000 Neuchâtel, Switzerland; Linguistic Research Infrastructure (LiRI), University of Zurich, 8050 Zurich, Switzerland; Department of Computational Linguistics, University of Zurich, 8050 Zurich, Switzerland; Zurich Forensic Science Institute (FOR), 8004 Zurich, Switzerland

**Keywords:** sexual dimorphism, geometric morphometrics, real-time MRI, articulatory variation

## Abstract

Perceptual experiments show that listeners recognize female speakers with lower accuracy than male speakers. Automatic speaker recognition systems may also show performance bias against female speakers even when training data sets are gender balanced. The underlying reasons for this discrepancy are unclear. Here, we apply geometric morphometrics to quantify sex-related morphological vocal tract disparity – the extent of shape variation – across both resting and articulatory configurations. We find that male speakers exhibit greater disparity in both resting and articulatory configurations. This morphological idiosyncrasy may in turn generate more discriminable acoustic signatures and offer a biological explanation for higher recognition accuracies for male voices by humans and machines. Our results suggest that innate variation in vocal tract morphology may contribute to performance bias in voice technology and voice perception by human listeners.

## 1. Introduction

Previous research has reported higher recognition accuracies for men in human and automatic recognition experiments [1, 2, 3, 4]. Speaker recognition systems are frequently trained on gender-unbalanced data containing more male speakers and therefore exhibit a performance bias against female speakers [5, 4, 6, 7]. Some studies show that even when the training data sets are balanced with respect to gender, voice recognition remains worse for female speakers [4]. In human perception experiments, male speakers are recognized with higher accuracy [1, 2, 3]. In addition to recognition discrepancy, a response bias is observed – when hearing younger and female speaker pairs, listeners are biased towards perceiving them as originating from the same speaker [3].

Different anatomical and sociolinguistic factors may underlie such recognition differences. Perception of speaker gender relies on a set of acoustic features interacting in a complex way, most notably voice fundamental frequency (f0), formant frequencies, and temporal information [8, 9]. Due to both physiological and cultural factors, male voices exhibit overall lower f0, resulting in denser spacing of harmonics compared to female voices [10, 11]. However, these differences are not universal and depend on language and cultural expectations of gender [12, 13]. As such, it is unclear whether this comparatively sparser harmonic sampling in female voices reduces the extent to which vocal tract idiosyncrasies are reflected in their spectral envelopes and whether this affects voice recognition by humans and automatic systems.

In this study, we apply geometric morphometrics (GM) [14] to quantify sex-related differences in resting state and articulatory vocal tract morphology, with a focus on morphological disparity, i.e., the extent of shape variation within and between sexes. Previous studies evaluated sexual dimorphism in vocal tract and articulatory patterns [15, 16, 10, 17, 18]. However, whether there is a relationship between patterns and magnitude of variation within and between sexes and voice identification has not been properly researched. Here we test whether greater morphological disparity in male speakers correlates with documented biases in voice recognition systems.

## 2. Methods

We analyzed resting state and articulatory vocal tract MRI data from 73 adult speakers (38 female, 35 male) drawn from the USC Speech and Vocal Tract Morphology MRI Database [19], including a subset of 32 speakers (16 female, 16 male) from the same cohort who produced the target vowels /i, e, a, o, u/ embedded in carrier words. Resting state morphology was quantified from midsagittal slices of the static volumetric MRI data, acquired while speakers were in a resting position. Articulatory vowel configurations were quantified from the 2D real-time MRI acquisitions. All MRI scans were acquired in the midsagittal plane, and vocal tract morphology was quantified by placing 2D anatomical landmarks on the image data. For the articulatory dataset, landmark configurations were first extracted along the dynamic articulatory trajectory of the vowel token of interest within each carrier word, rather than across the whole carrier word. Multiple frames were averaged across the two analyzed repetitions, yielding a single representative shape per speaker and vowel. We employed two distinct landmarking protocols: one optimized for resting state vocal tract configurations (Fig. 1A) and another tailored for articulatory vowel data (Fig. 2A). Landmarks were placed using the software package StereoMorph in *R* [20, 21, 22].

**Figure 1.**
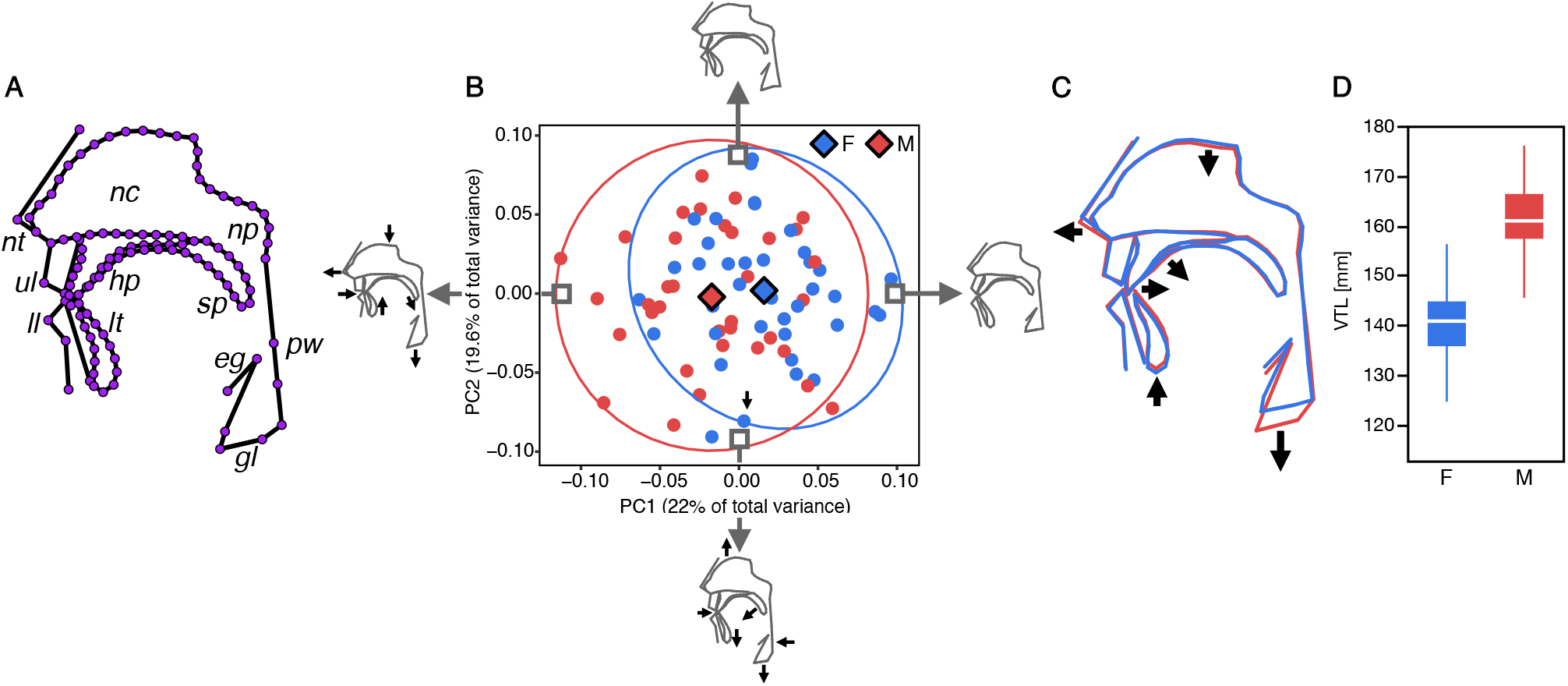
Resting state vocal tract variation between sexes. **A:** Anatomical landmarks used in the analysis of resting state vocal tract MRI data. Abbreviations: nt: nasal tip, nc: nasal cavity, np: nasopharynx, ul: upper lip, hp: hard palate, sp: soft palate, pw: pharyngeal wall, ll: lower lip, lt: lower teeth, eg: epiglottis, gl: glottis. **B:** PCA of resting state vocal tract shapes. Data points are colored by sex (blue: female, red: male), with diamond markers indicating group means and 90% density ellipses outlining within-sex shape variation. Shape changes along major PC axes are illustrated by gray configurations at the extreme ends of variation, with black arrows indicating the direction of anatomical change associated with each axis. **C:** Average resting state vocal tract shape by sex: female (blue), male (red). Black arrows indicate major shape changes from female to male configurations. **D:** Boxplot of vocal tract length differences between sexes (mm).

**Figure 2.**
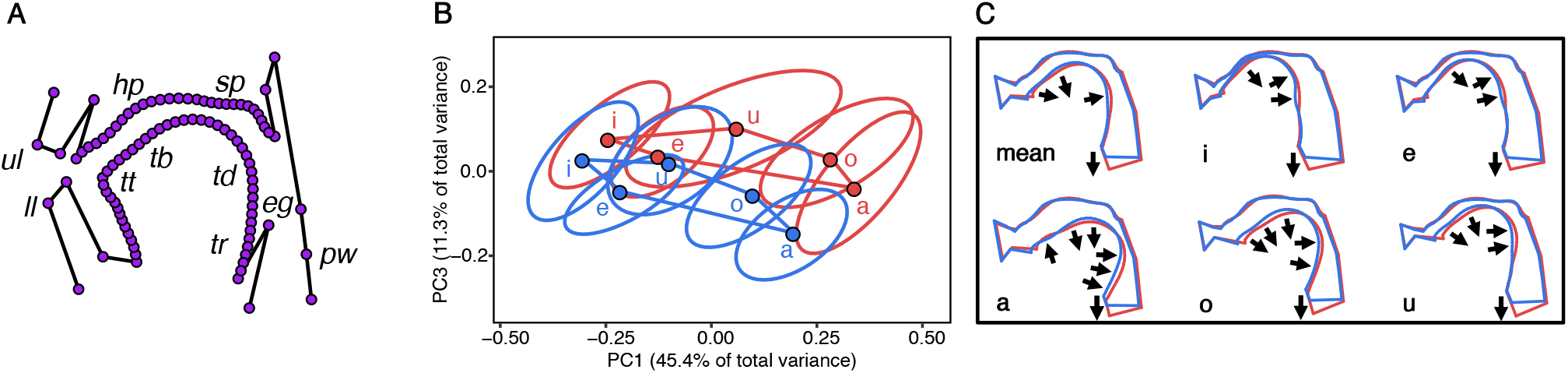
Articulatory shape space by sex. **A:** Anatomical landmarks used in the analysis of dynamic vowel MRI data. Abbreviations: tt: tongue tip, tb: tongue blade, td: tongue dorsum, tr: tongue root, for remaining abbreviations see Fig. 1. **B:** PCA of dynamic vocal tract shapes, shown separately for each vowel. Data are colored by sex (blue: female, red: male), with dots indicating vowel- and sex-specific means. 90% density ellipses outline within-vowel, within-sex shape variation. Vowel means are connected to form a vowel polygon, illustrating the articulatory vowel space. We visualize PC1 (primarily vowel-related variation) against PC3 (capturing sex- and vowel-related variation), excluding PC2 (primarily interindividual variation) to visually focus on vowel structure and between-sex differences. **C:** Average articulatory shape by sex (female: blue, male: red), shown per vowel and for the pooled mean. Black arrows indicate major shape changes from female to male configurations.

To quantify shape variation in resting state and articulatory vocal tract configurations, as well as to assess overall vocal tract size, we applied geometric morphometrics (GM) [14]. Digitized landmark configurations were aligned using Generalized Procrustes Analysis (GPA) to remove non-shape variation, i.e., differences in position, orientation, and scale [23]. Principal component analysis (PCA) was subsequently performed on the Procrustes-aligned coordinates to visualize and interpret multivariate shape variation (Figs. 1B & 2B). All GM analyses were conducted in *R* using the packages geomorph and Morpho [22, 24].

Vocal tract size was quantified using centroid size (Csize) of the resting state landmark configurations, defined as the square root of the sum of squared Euclidean distances from each landmark to the configuration centroid. Centroid size provides a measure of overall size and is widely used in morphometric analyses [25, 26, 14].

Vocal tract length was measured manually on resting state vocal tract images in *ImageJ* [27] as the curvilinear distance along the vocal-tract midline from the anterior lips, through the oral and pharyngeal cavities, to the glottis, following previous MRI-based approaches to vocal-tract length measurement [15, 28].

Morphological disparity, defined as within-group Procrustes variance, was quantified as the mean squared distance of individual shapes from their group mean in Procrustes shape space (i.e., after GPA-alignment of landmarks). Disparity was computed for resting state and articulatory datasets using the *morphol*.*disparity* function in geomorph [22]. Disparity was estimated separately for each sex and vowel category, with centroid size included as a covariate to account for potential allometric effects, i.e., residual size-related shape variation [26]. In addition, for the articulatory dataset we evaluated disparity in a pooled analysis across vowels, including vowel category as an additional covariate in the model. Sex differences in disparity were statistically assessed using permutation tests (*n*=999) under the null hypothesis of no group difference. In each iteration, residual Procrustes coordinates (after accounting for centroid size) were randomly reassigned across groups to generate a null distribution of disparity differences.

Procrustes variances are reported in Fig. 3A. To localize anatomical sources of disparity, we calculated per-landmark variance differences for both resting state and articulatory datasets. For each landmark, variance was computed as the summed variance across the two coordinate dimensions of the Procrustes-aligned data. Female variance was subtracted from male variance to obtain sex-specific disparity differences. These differences were visualized as heatmaps overlaid on the mean vocal tract configurations (Figs. 3B–C), where color encodes the direction and magnitude of disparity differences: blue/purple indicates greater female variability, white indicates no difference, and green/yellow indicates greater male variability. This visualization enables spatial localization of sex-related differences in morphological and articulatory variability.

**Figure 3.**
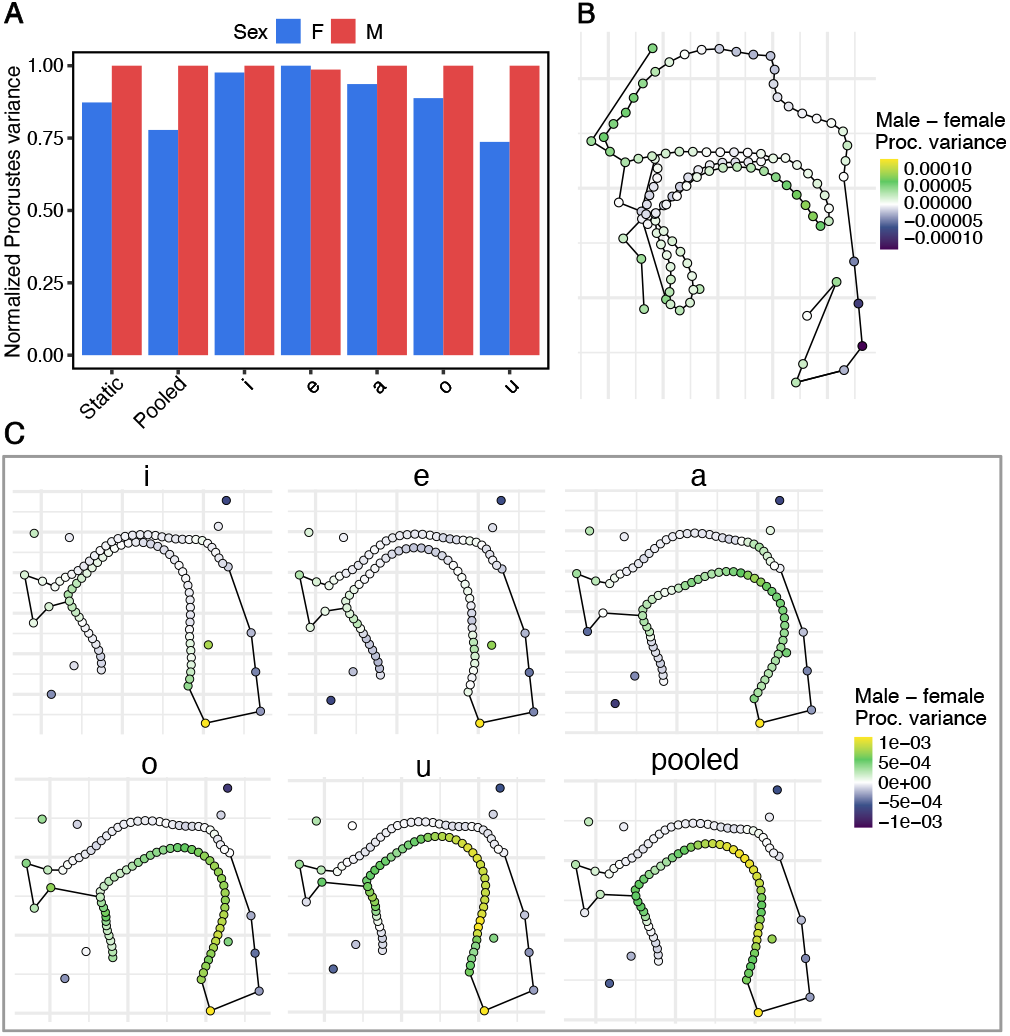
Sex differences in morphological disparity. **A:** Normalized Procrustes variance (morphological disparity) for resting state vocal tract data, pooled articulatory data, and per vowel /i e a o u/. Values are scaled to maximum within each dataset (max = 1). **B:** Male–female disparity per landmark of resting state vocal tract shape. Each point represents a vocal tract landmark, colored according to the difference in Procrustes variance between male and female speakers (male – female). Blue/purple indicates greater female variability, white indicates no difference, and green/yellow indicates greater male variability. **C:** Male–female disparity per landmark in articulatory data. Heatmaps show the difference in Procrustes variance (male – female) for each vocal tract landmark during articulation of vowels /a/, /e/, /i/, /o/, /u/, and across all vowels (pooled). Color scheme same as in **B**.

## 3. Results

Figure 1B illustrates resting state vocal tract shape variation across sexes. Between-sex differences are primarily structured along PC1, which reflects variation in laryngeal position and variation in the anteroposterior position of the maxillomandibular complex (prognathism vs. retrognathism). Mean shape comparisons indicate that the most pronounced sex difference is the lower larynx position in males (Fig. 1C), corresponding to substantially greater vocal tract length relative to females (Fig. 1D). In addition to shape differences, males exhibit larger overall vocal tract size, with centroid size averaging 1503.04 in males compared to 1386.16 in females, corresponding to an approximately 8.4% increase in males. Males also exhibit greater morphological disparity than females in the first two principal components (PC1–PC2), which together account for 41.6% of total shape variation in the sample. This increased dispersion is evident from the larger density ellipses in PC1–PC2 shape space (Fig. 1B), indicating elevated within-sex shape variability in males. Variation along PC2, i.e., within-sex variation, primarily contrasts relative oral and pharyngeal cavity length: individuals at one end of the distribution display greater anteroposterior extension of the oral cavity, whereas those at the opposite end exhibit superoinferior elongation of the pharyngeal cavity.

Figure 2B illustrates articulatory shape variation between male and female speakers for each vowel. In both sexes, articulatory configurations form distinct vowel-specific polygons in PC1–PC3 shape space (while PC2 primarily captures non-sex-related interindividual variation in the sample), reflecting the characteristic vocal tract postures associated with /i, e, a, o, u/. However, males consistently exhibit greater articulatory excursions across most vowels, particularly for the back vowels (/a/, /o/, /u/), as evidenced by the larger spatial extent of their density ellipses relative to those of females, as well as by a larger average vowel polygon (i.e., greater inter-vowel distances). This broader dispersion indicates a greater range of articulatory configurations, i.e., more morphological disparity in males during vowel production. Part of this increased excursion may relate to generally larger and longer vocal tract dimensions in males.

Comparisons of mean articulatory shapes further show that males tend to occupy more extreme configurations, especially for back vowels (Fig. 2C). These configurations are characterized by enhanced posterior and lower tongue positions, a pattern that may be associated with the typically lower laryngeal position in males, which could permit greater superoinferior and posterior shaping of the pharyngeal cavity.

Beyond the visual patterns evident in multivariate shape space, a formal statistical evaluation of morphological disparity in the pooled articulatory dataset reveals that males exhibit significantly greater shape variation than females during articulation (p = 0.006; Fig. 3A), supporting the hypothesis that male vocal tracts are more variable in their movement during speech. In contrast, no significant sex differences in morphological disparity emerged in resting state vocal tract morphology or in vowel-specific configurations. This may reflect reduced statistical power in smaller per-vowel samples, as well as greater anatomical constraint in resting postures, where sex differences in variability are comparatively subtle (cf. Figs. 1B–C). Although non-significant, the trend across all datasets – with the exception of /e/ – consistently favors higher male disparity (Fig. 3A), suggesting that the effect becomes pronounced under the integrated articulatory demands of vowel production.

To elucidate the anatomical basis of this disparity, we mapped per-landmark variance differences onto the vocal tract geometry (Fig. 3B—C). In resting state configurations, males exhibit greater variability across multiple regions, notably the anterior nasal cavity, mandibular symphysis (chin), lower dental arch, and anterior laryngeal complex (glottis and epiglottis). Females, by contrast, show relatively elevated variation in the maxillary arch (hard palate and anterior teeth) and the posterior pharyngeal wall.

During articulation, the disparity becomes more spatially concentrated and functionally structured. In the pooled analysis across vowels (Fig. 3C), males display significantly greater variability in the laryngeal root (epiglottis) and across the tongue body, particularly the dorsum, consistent with a broader range of tongue positioning and laryngeal adjustment. This effect is especially pronounced for back vowels (/a/, /o/, /u/). For front vowels (/i/, /e/), males additionally exhibit heightened variability at the tongue tip and lip landmarks. Females, in turn, show increased articulatory variability in the tongue dorsum during production of front vowels and consistently across all vowels at the chin tip, indicating relatively greater jaw-related variation. As in resting state morphology, female speakers also exhibit elevated variation in the posterior pharyngeal wall and hard palate.

## 4. Discussion

Our results show that males exhibit greater vocal tract variation than females. This suggests that male vocal tracts are more idiosyncratic in structure and function, which may be one of the factors contributing to better recognition of male speakers by human listeners [1, 2, 3] and by automatic systems trained on gender-balanced data [4]. Therefore, it is plausible that greater morphological variability in male vocal tracts produces more distinct acoustic signatures. This, in turn, could contribute to greater perceived distinctiveness in male voices, offering a plausible anatomical and acoustic explanation for higher recognition accuracies for male speakers. At the same time, given that human listeners and automatic systems rely on partly different acoustic cues for voice processing [29], future research should systematically examine how vocal tract disparity relates to recognition performance across both human listeners and automatic systems.

From a technological perspective, our results have implications for fair and inclusive automatic voice recognition design and feature extraction methods (cf. [30]). Greater structural dispersion in male voices could mean that current feature extraction methods are more optimized for capturing speaker-specific details in male speech. If state-of-the-art deep neural network based automatic speaker recognition systems (e.g., ResNet [31] and ECAPA-TDNN [32]) exhibit systematic performance degradation for female speakers when trained on gender-balanced data, this may indicate that some performance discrepancies can be attributed to how effectively models encode speaker-specific acoustic properties, rather than biases in training data per se. Systematic investigation of the effects of harmonic density manipulations on automatic speaker recognition performance, as well as how speakers are clustered in multidimensional embedding space, are essential for elucidating and mitigating gender biases in current voice technologies.

The heightened vocal tract variability and articulatory variability in males is most prominent in back vowels and also shows higher laryngeal movement, indicating possibly more laryngeal adjustment during speaking, something that could explain the more variable positioning of the tongue in back vowels. These results align with findings from the forensic phonetics domain suggesting that low and back vowels exhibit wider variation in acoustic space across speakers compared to high front vowels [33]. While our analysis focuses on vowels, future work should extend these findings to fluent speech and naturalistic utterances to further test whether this morphological bias translates into broader performance asymmetries across speech domains.

The increased morphological disparity in males may also reflect an evolutionary adaptation linked to sexual selection. In human evolutionary history, male vocal traits – including their pitch, timbre, and resonance – possibly served dual roles: attracting mates and asserting dominance over rivals [34, 35, 36, 37]. Functions requiring reliable individual recognition (e.g., coalition formation or competitive interactions) would likely benefit from idiosyncratic acoustic cues that differentiate individuals [38, 39, 40, 41]. Greater morphological and acoustic variability among males, potentially arising from sexual selection on dominance and formidability, may therefore contribute to increased vocal discriminability as a secondary consequence. By contrast, female vocal traits are more consistently linked in the literature to perceptions of youth and attractiveness [42, 43, 44], suggesting distinct selective pressures across sexes [45].

More broadly, our findings highlight the importance of comprehensive approaches that jointly examine speakers’ anatomy, articulation, and acoustics, and their effects on voice and speech recognition. While male voices have been associated with higher identity recognition, female speakers are frequently reported as more intelligible and clearer speakers, due in part to larger vowel spaces and more precise articulation [46, 47, 48, 11, 49]. This dissociation may suggest that voice recognizability (who is speaking?) and speech intelligibility (what is being said?) may rely on distinct acoustic, articulatory, and anatomical mechanisms. Elucidating these mechanisms is crucial for advancing speech and voice technology, as well as improving theoretical models of speech communication.

## 5. Acknowledgments

Volker Dellwo provided key insights into the relationship between vocal tract idiosyncrasy and speaker recognizability. This work was funded by the Swiss National Science Foundation (PCEFP1 186841, EVOPHON).

## 6. Generative AI Use Disclosure

Generative AI tools were used to assist with language editing for improving clarity of the manuscript text.

## Notes

### Competing Interest Statement

The authors have declared no competing interest.

